# Identification of a novel tetrameric structure for human apolipoprotein-D

**DOI:** 10.1101/335232

**Authors:** Claudia S. Kielkopf, Jason K.K. Low, Yee-Foong Mok, Surabhi Bhatia, Tony Palasovski, Aaron J. Oakley, Andrew E. Whitten, Brett Garner, Simon H.J. Brown

**Affiliations:** Illawarra Health and Medical Research Institute, University of Wollongong, Wollongong, NSW, Australia; School of Biological Sciences, University of Wollongong, Wollongong, NSW, Australia; Molecular Horizons, University of Wollongong, Wollongong, NSW, Australia; School of Life and Environmental Sciences, University of Sydney, Sydney, NSW, Australia; Department of Biochemistry and Molecular Biology, Bio21 Molecular Science and Biotechnology Institute, University of Melbourne, Parkville, VIC, Australia; Illawarra and Shoalhaven Local Health District (ISLHD), Wollongong, NSW, Australia; Specialist Breast Clinic Sutherland Shire and Wollongong, NSW, Australia; Integrated Specialist Health Care Sutherland Shire, NSW, Australia; School of Chemistry, University of Wollongong, Wollongong, NSW, Australia; Australian Nuclear Science and Technology Organisation, Lucas Heights, NSW, Australia

**Keywords:** Lipocalin, oligomerization, apolipoprotein structure, small-angle x-ray scattering (SAXS), lipid, lipocalin structure

## Abstract

Apolipoprotein-D is a 25 kDa glycosylated member of the lipocalin family that folds into an eight-stranded β-barrel with a single adjacent α-helix. Apolipoprotein-D specifically binds a range of small hydrophobic ligands such as progesterone and arachidonic acid and has an antioxidant function that is in part due to the reduction of peroxidised lipids by methionine-93. Therefore, apolipoprotein-D plays multiple roles throughout the body and is protective in Alzheimer’s disease, where apolipoprotein-D overexpression reduces the amyloid-β burden in Alzheimer’s disease mouse models.

Oligomerisation is a common feature of lipocalins that can influence ligand binding. The native structure of apolipoprotein-D, however, has not been conclusively defined. Apolipoprotein-D is generally described as a monomeric protein, although it dimerises when reducing peroxidised lipids.

Here, we investigated the native structure of apolipoprotein-D derived from plasma, breast cyst fluid (BCF) and cerebrospinal fluid. In plasma and cerebrospinal fluid, apolipoprotein-D was present in high-molecular weight complexes, potentially in association with lipoproteins. In contrast, apolipoprotein-D in BCF formed distinct oligomeric species. We assessed apolipoprotein-D oligomerisation using native apolipoprotein-D purified from BCF and a suite of complementary methods, including multi-angle laser light scattering, analytical ultracentrifugation and small-angle X-ray scattering. Our analyses showed that apolipoprotein-D predominantly forms a ∽95 to ∽100 kDa tetramer. Small-angle X-ray scattering analysis confirmed these findings and provided a structural model for apolipoprotein-D tetramer. These data indicate apolipoprotein-D rarely exists as a free monomer under physiological conditions and provide insights into novel native structures of apolipoprotein-D and into oligomerisation behaviour in the lipocalin family.

## INTRODUCTION

Human apolipoprotein-D (apoD) is a 169 amino acid glycoprotein member of the lipocalin family. In this protein family, affiliation is not based on sequence homology but rather on structural homology (Akerstrom et al., 2000). The crystal structure of an apoD monomer reveals a typical lipocalin fold with an eight-stranded antiparallel β-barrel flanked by an α-helix (Eichinger et al., 2007). Two disulphide bridges tether loops to the central β-barrel and a fifth cysteine is free (Eichinger et al., 2007). The apoD β-barrel encloses a conical shaped hydrophobic cavity, referred to as the apoD ligand-binding pocket. Early studies suggested that apoD can bind a range of lipids, including arachidonic acid (AA), cholesterol and several steroids (Dilley et al., 1990; Lea, 1988; Morais Cabral et al., 1995; Pearlman et al., 1973). More recent studies indicate that the binding of lipids in the apoD binding pocket is in fact specific (Eichinger et al., 2007; Vogt and Skerra, 2001). Progesterone and AA both bind to the apoD binding pocket with high affinity, whereas pregnenolone and specific eicosanoids (e.g. 12-HETE and 5,15-diHETE) bind with reduced affinity (Dilley et al., 1990; Lea, 1988; Morais Cabral et al., 1995). Cholesterol does not appreciably bind within the apoD binding pocket (Morais Cabral et al., 1995). Additionally, apoD may also interact with lipids via a region of surface hydrophobicity, due to exposed hydrophobic residues located in three of its extended loops (Eichinger et al., 2007). Located close to the open end of the binding pocket, this hydrophobic cluster may facilitate apoD association with high-density lipoprotein (HDL) particles and permit insertion of apoD into cellular lipid membranes (Eichinger et al., 2007). This hydrophobic surface explains observations that apoD binds a range of lipophilic molecules. We have shown that oxidised (hydroperoxy) forms of eicosatetraenoic acids bind to this hydrophobic surface, where the potential radical-generating lipid hydroperoxide (L-OOH) moiety is reduced to an inert lipid hydroxide (L-OH), thereby inhibiting lipid peroxyl radical “propagation” of free radical-mediated lipid oxidation (Bhatia et al., 2012a; Oakley et al., 2012). This antioxidant action of apoD is critically dependent on the redox active side chain of M93 that is reversibly converted to methionine sulfoxide (MetSO) in the reaction. Creation of the MetSO destabilises an extended loop close to the entrance of the ligand binding pocket which in turn promotes apoD dimerisation (Bhatia et al., 2012a; Oakley et al., 2012).

ApoD is cleaved from a 189 amino acid precursor and has two N-linked glycosylation sites on N45 and N78 that in plasma are mainly trisialo triantennary (N45) and fucose disialo biantennary (N78) structures (Schindler et al., 1995). For consistency, all residue numbers given in this manuscript are based on the mature apoD sequence. The addition of the theoretical glycan masses to the molecular weight of apoD leads to a total mass of 24.52 kDa. However, on reducing SDS-PAGE, apoD is retained at an apparent molecular weight of up to 32 kDa (Balbín et al., 1990). The physiological range of apoD concentration depends on the fluid. The apoD concentration in plasma ranges from 0.05 to 0.2 mg/ml (Van Dijk et al., 2013), in breast cyst fluid (BCF) ranges from 13.7 to 15.1 mg/ml (Sánchez et al., 1992) and in cerebrospinal fluid (CSF) is reported to be 0.0012 mg/ml (Terrisse et al., 1998). The glycosylation pattern of apoD depends on the expression site, as apoD glycosylation in axillary secretion (Zeng et al., 1996) differs from plasma (Schindler et al., 1995), which in turn is different from apoD expressed in the brain (Li et al., 2016).

ApoD appears to play a protective role in Alzheimer’s disease (AD) most likely by combating oxidative stress and neuroinflammation. Increased apoD dimer formation is observed in AD brain samples (Bhatia et al., 2013), suggesting that apoD can directly abate oxidative stress in AD by reducing peroxidised lipids. Accordingly, apoD expression is increased in the brain and CSF of AD patients (Terrisse et al., 1998). In mouse studies, overexpression of apoD in an AD model decreases the Aβ burden in the brain (Li et al., 2015). Furthermore, apoD was recently shown to protect a vulnerable subset of lysosomes under oxidative stress conditions (Pascua-Maestro et al., 2017).

Previous studies show that the formation of dimers or higher order oligomers is a common feature in lipocalins (Akerstrom et al., 2000; Gasymov et al., 2007; Huber et al., 1987a). Such oligomerisation has been characterised by size exclusion chromatography (SEC) and multi-angle laser light scattering (MALLS) (Gasymov et al., 2007), analytical ultracentrifugation (AUC) (Gouveia and Tiffany, 2005), small-angle X-ray scattering (SAXS) and native PAGE (Kozak and Grubb, 2007). Interestingly, the oligomeric state of lipocalins has been linked to their ligand binding functions (Gamiz-Hernandez et al., 2015; Gutierrez-Magdaleno et al., 2013). Ligand binding of β-lactoglobulin leads to dimer dissociation (Gutierrez-Magdaleno et al., 2013) and the function of crustacyanin as pigmentation protein is critically dependent on dimer formation (Gamiz-Hernandez et al., 2015). Unlike these lipocalins, apoD is described as a monomer by most publications (Akerstrom et al., 2000; Nasreen et al., 2006), apart from reports of non-disulphide linked dimers formed upon reducing peroxidised lipids *in vitro* (Bhatia et al., 2012a) and in Alzheimer’s disease (AD) brain tissue (Bhatia et al., 2013). Disulphide-linked apoD dimers are found in urine (Blanco-Vaca and Pownall, 1993) and tears (Holzfeind et al., 1995).

Given that oligomerisation is a common feature of other lipocalins, in the present study we investigated the native structure of apoD in human apoD-containing fluids. Specifically, utilising BCF as source of apoD, we examined the native apoD structure in a suite of complementary analytical experiments that enabled us to generate a robust data set which, for the first time, indicate that apoD forms a tetramer under native conditions.

## MATERIALS AND METHODS

### Chemicals and Reagents

All chemicals were biotechnology grade or similar. Tris, piperazine and glycerol were sourced from Sigma (Castle Hill, NSW, Australia), NaCl and ammonium sulphate were from Astral Scientific (Gymea, NSW, Australia), mono-and di-sodium phosphate from Ajax Finechem and Sigma (Castle Hill, NSW, Australia), respectively, and DTT from Astral Scientific (Gymea, NSW, Australia).

### Fluid sample collection and preparation

Breast examination was performed and BCF was collected by point of care ultrasound with aspiration of cyst by TP. BCF was stored at 4°C and transferred from the clinic to the laboratory within 4 hours of collection. A protease inhibitor cocktail (Sigma, 1:100 dilution) was added, particulate was removed by centrifugation at 23k × *g* for 20 min and BCF aliquots were stored at -80°C.

Plasma was obtained by collection of 5 ml of whole blood in EDTA collection tubes (BD vacutainer), incubated at 22°C for 30 min and then centrifuged for 10 min at 1400 × *g*. The plasma fraction was carefully transferred to a new tube without disturbing the red blood cells. Protease inhibitor cocktail (Sigma, 1:100 dilution) was added and plasma aliquots were stored at -80°C.

CSF (Lee Bioscience) consisted of pooled CSF from healthy human subjects. The obtained vials were stored at -80°C and before use, thawed on ice and protease inhibitor cocktail (Sigma, 1:100 dilution) was added.

All fluids were thawed on ice and particulates were removed by centrifugation at 16k × *g* for 20 min at 4°C immediately before use. An overview of the following experimental design is shown in Figure 1.

**Fig. 1.**
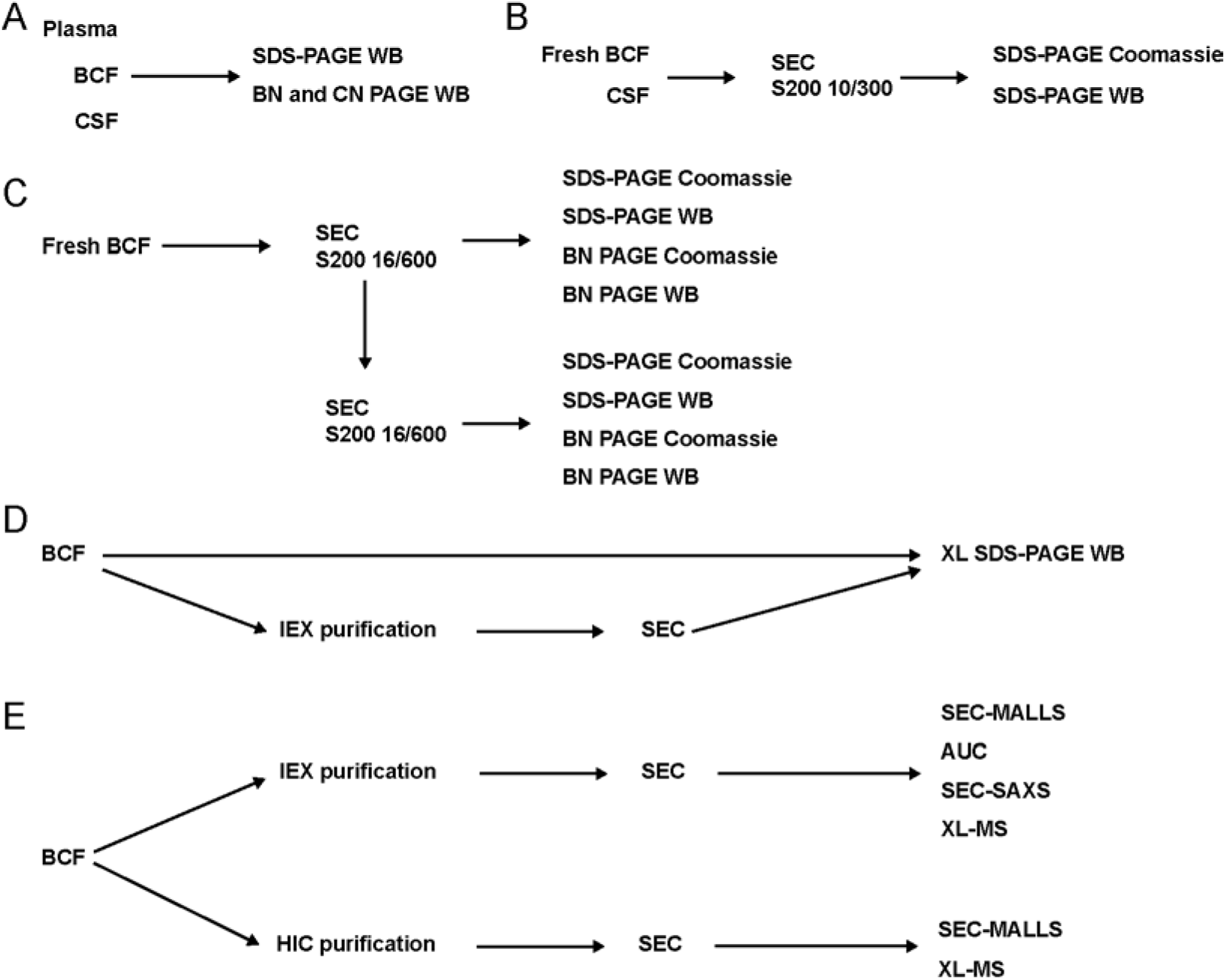
Overview of the experimental design. A) - E) outline the experimental process and summarise the fluids and purified apoD samples used for procedures in this study.

### Characterisation of BCF apoD by FPLC and HPLC SEC

FPLC was performed on an NGC Scout system (Bio-Rad) and HPLC on an 1100 Series HPLC system (Agilent).

BCF was applied neat or at the dilution described below to three formats of SEC; GE Superdex 200 16/600 (bed volume ∽120 ml), GE Superdex 200 Increase 10/300 (bed volume ∽24 ml), and a TOSOH TSKgel Ultra SW Aggregate (bed volume ∽14 ml). Fraction volumes collected were 2 ml (S200 16/600) and 5 μl (S200 10/300). Superdex columns were equilibrated into buffer containing 25 mM Tris, 150 mM NaCl, pH 8.0, using the Superdex 16/600 at a flow rate of 0.75 ml/min with a BCF injection volume of 200 μl, and the Superdex 10/300 at a flow rate of 0.5 ml/min with a BCF injection volume of 100 μl (diluted 1:1 with buffer). TOSOH TSKgel Ultra SW Aggregate chromatography was performed using a buffer containing 100 mM sodium phosphate, 100 mM sodium sulphate at pH 6.7 at a flow rate of 1.0 ml/min, and all samples (10 μl, BCF 1:20 dilution) were injected in triplicate.

### Characterisation of CSF apoD by FPLC SEC

CSF (250 μl) was applied neat to the GE Superdex 200 Increase 10/300 column, which was equilibrated with 25 mM Tris, 150 mM NaCl, pH 8.0, at a flow-rate of 0.5 ml/min.

### Calibration of SEC columns and mass calculations

SEC columns were equilibrated into the appropriate buffers (see above) and calibrated with an injection (Superdex 200: 250 μl, TOSOH: 10 μl) of SEC protein standard mix (Sigma) containing bovine thyroglobulin (670 kDa), bovine γ-globulins (150 kDa), chicken egg albumin (44.3 kDa) and bovine pancreas ribonuclease (13.7 kDa). The standard and sample peaks were integrated to determine the retention volume using ChromLab software (BioRad) or ChemStation (Agilent). The partition coefficient K_av_ was calculated via the bed height and void volume of each column according to equation 1:

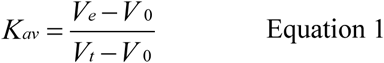

with V_e_: Elution volume, V_o_: Void volume, V_t_: total column volume. K_av_ and Log_10_ of the molecular weight in Da of each standard was plotted and fitted with a linear regression curve. Calculated error ranges were plotted. Using this standard curve, the molecular weights of the unknown samples were calculated.

### Protein Purification

BCF fractions collected from the Superdex 200 16/600 SEC were characterised by SDS-PAGE, followed by Coomassie staining (G-250, SimplyBlue™ SafeStain) and western blotting. Fractions highly enriched in apoD (1 ml of each fraction covering eluate from 64.4 ml to 70.4 ml) were pooled, concentrated using Amicon Ultra concentrators with an Ultracel-10 membrane (10,000 Da molecular weight cut-off), and applied to the Superdex 200 10/300.

Hydrophobic interaction chromatography (HIC) purification of apoD was performed on a GE HiScreen Butyl-S FF column equilibrated into HIC buffer (50 mM sodium phosphate, 1.5 M ammonium sulphate, pH 7.0). BCF aliquots of 250 μL were diluted into 15 ml of HIC buffer, applied to the column and then eluted with a 0 - 100% linear gradient using ultrapure water over 4 column volumes (CV).

Ion exchange (IEX) chromatography purification of apoD was performed on a GE HiTrap ANX™ Sepharose FF 5 ml column equilibrated into IEX buffer (20mM piperazine, pH 5.0). BCF aliquots of 250 μL were applied to the column and eluted with a 0-50% gradient using high salt buffer (20 mM piperazine, 1 M NaCl, pH 5.0) over 4 CV.

IEX or HIC purified apoD fractions were identified by Coomassie SDS-PAGE, pooled, concentrated and buffer exchanged to 25 mM Tris, 150 mM NaCl, pH 8.0. Subsequently, apoD was purified in a polishing SEC step using the Superdex 200 16/600 column. Protein concentrations were measured using a Nanodrop 2000c (ThermoFisher, Abs 280 nm, ε=32,680) or a Pierce BCA assay (ThermoFisher) with bovine serum albumin (BSA) serial dilution as standard curve according to the manufacturer’s instructions.

### SDS-PAGE and western blotting

Samples for SDS-PAGE were prepared by adding 4× sample buffer (Invitrogen) and DTT to a final concentration of 5 μM, denatured for 10 min at 70°C and separated on 4-12% BOLT Bis-Tris SDS-polyacrylamide gels using MES running buffer (both Invitrogen) for 22 min at 200 V at 22°C. The gels were either stained using Coomassie staining (G-250, SimplyBlue™ SafeStain) or used for western blotting and immunodetection. SDS gels were either incubated in 20% (v/v) ethanol for 10 min and then transferred to 0.4 μm PVDF membranes using the iBLOT semi-dry blotter (7 min programme P0) or transferred to 0.4 μm PVDF for 1 h at 20 V using mini Blot modules (all Invitrogen). Post-transfer, the membranes were rinsed in reverse osmosis H_2_O, stained using Ponceau S (0.1% (w/v) in 5% acetic acid, Sigma) and then incubated in hot PBS for 10 min as an antigen-retrieval step. After blocking in PBS-10% Tween-20 (PBS-T) with 5% (w/v) skim milk powder for >1 h at 22°C, the membranes were probed for human apoD with monoclonal mouse anti-human apoD C1 antibody (1:5,000, Santa Cruz, SC-373965, lot G1911) in PBS-T with 5% (w/v) skim milk at 4°C overnight. Membranes were washed in PBS-T (1 min, then 3×5 min) and then incubated for 2 h at 22°C with polyclonal goat anti-mouse IgG conjugated to horseradish peroxidase (1:5,000, Dako, P0447, lot 00077665). After washing in PBS-T, the chemiluminescence signal was detected using Pierce ECL plus substrate (ThermoFisher) and a CCD imager (Amersham Imager 600RGB).

### Blue and clear native PAGE

BisTris NativePAGE running buffer and Tris-Glycine native buffer (Invitrogen) were prepared according to the manufacturer’s instructions and 1% (v/v) NativePAGE cathode additive was added to BisTris NativePAGE buffer. Buffers were then chilled to 4°C. Samples were prepared by adding sample buffer to 1 × and then separated on 4-16% NativePAGE Bis-Tris gels (Invitrogen) for 90 min at 150 V or on 4-12% Novex Tris-Glycine gels (Invitrogen) for 60 min at 125 V, on ice. Gels were either stained using Coomassie staining (0.02% R-250 (w/v) in 10% (v/v) acetic acid, 40% (v/v) methanol, then destained in 8% (v/v) acetic acid) or used for western blotting. Native gels were incubated in 2× transfer buffer with 10% methanol for 10 min and then transferred to 0.4 μm PVDF membranes using the iBLOT semi-dry blotter (7 min programme P0). Post-transfer, the membranes were incubated for 15 min in 8% acetic acid, rinsed in reverse osmosis H_2_O and then incubated in hot PBS for 10 min for antigen-retrieval. Membranes were dried, reactivated in methanol and stained using Ponceau S. The immunodetection procedure was the same as described for SDS-PAGE but the apoD C1 antibody was used at a 1:1,000 dilution.

### Crosslinking

A 2 mg BS^3^ aliquot (bis(sulfosuccinimidyl)suberate, ThermoFisher) was equilibrated to 22°C and dissolved immediately before use in ultrapure water to a concentration of 50 mM. Crosslinking was performed on BCF and IEX-purified apoD tetramer in parallel, at a final apoD concentration of 10 μM and final volume of 200 μl (BCF) or 50 μl (purified apoD). Samples were prepared in conjugation buffer (100 mM sodium phosphate, 150 mM NaCl, pH 7) and BS^3^ was added at 10 ×, 20 ×, 50 ×, 100 × and 200 × molar excess. Samples were mixed 30 s in vortex mixer, centrifuged at 1000 × g for 1 min and incubated at 22°C for 30 min or 1 h. The reaction was quenched for 15 min by adding quenching buffer (1M Tris-HCl, pH 7.5) to a final concentration of 50 mM Tris, vortexed and spun. Appropriate sample volumes were removed, mixed with SDS-PAGE loading dye and reducing agent and analysed using SDS-PAGE western blot as described above. The bands intensity was quantified using ImageJ.

### Analytical ultracentrifugation

Sedimentation velocity experiments were performed on a Beckman Coulter XL-I analytical ultracentrifuge equipped with an An-Ti60 rotor; 400 μl of 0.08, 0.26, 0.4, 0.6 and 0.8 mg/ml HIC-purified apoD tetramer in 25 mM Tris with 150 mM NaCl at pH 8.0 were applied to the sample compartment of a double-sector centrepiece, with buffer in the reference compartment. The samples were centrifuged at 50,000 rpm for 10 h at 20°C and protein sedimentation was detected using absorbance at 280 nm. Data analysis was conducted using SedFit (Schuck and Rossmanith, 2000) by fitting the data to a continuous sedimentation coefficient model [c(s)]. The resulting size distribution curves were then normalised to enable comparison of the curves at different apoD concentrations.

### Multi-angle laser light scattering

HIC-purified apoD (100 μl, 2 mg/ml) or IEX-purified apoD (80 μl, 7 mg/ml) were analysed using SEC (S200 10/300 on an Äkta system, equilibrated with 25 mM Tris with 150 mM NaCl at pH 8.0) with online MALLS, UV absorbance and refractive index detectors (Wyatt). UV, MALLS, and dRI data were collected and analysed using ASTRA™ software (Wyatt Technology), and molecular weight determinations were carried out according to the Debye-Zimm model using a dn/dc value of 0.1876 ml/g for apoD (calculated by SedFit based on primary sequence). A run with BSA (100 μl, 2 mg/ml) was used to align UV, light scattering and refractive index signal.

### Crosslinking mass spectrometry (XL-MS)

For detailed method description, see Supporting Information Methods. Briefly, for each crosslinking experiment, 17-50 μg of IEX-and HIC-purified ApoD at ∽0.4-1.2 mg/mL was crosslinked using BS^3^ and adipic acid dihydrazide (ADH) and 4-(4,6-dimethoxy-1,3,5-triazin-2-yl)-4-methylmorpholinium chloride (DMTMM). DMTMM crosslinking is a side-reaction in the ADH crosslinking experiment. Samples were then digested with trypsin. For the generation of the MS/MS search database, 2-3 μg of purified, non-crosslinked apoD was also prepared for LC-MS/MS using trypsin. Peptides were then separated using a C18AQ particles column (Dr Maisch GmbH HPLC) on a Dionex UltiMate 3000 UHPLC system (ThermoFisher Scientific). Mass analyses were performed using a Q-Exactive Plus mass spectrometer (ThermoFisher Scientific) and identified using MSconvert tool (Chambers et al., 2012) and the database search program Mascot (Matrix science). For the screening and evaluation of potential crosslinks, a default FDR of 5% was used and only peptides with scores ≤1×10^-4^ (Yang et al., 2012) were considered for further analyses. All spectra that met these criteria were also manually visually verified: only crosslinks with at least four fragment ions on both the alpha-and beta-chain peptide each and addressing the most abundant peaks in the spectrum were retained for further analyses. Crosslinks that fulfilled all these conditions were deemed unambiguous and high-confidence. Theoretical and experimental masses, peptide sequences and how many times a specific crosslink was detected can be found in Table S2.

### Small angle X-ray scattering data collection and analysis

SEC-SAXS data in co-flow mode was collected at the SAXS/WAXS beamline at the Australian Synchrotron, Melbourne, Australia. IEX-and SEC-purified apoD in SAXS-buffer (50 mM Na Phosphate, 150 mM NaCl, 3% (v/v) glycerol, pH 7.4) was spin-concentrated (7.5 mg/ml, measured by BCA assay) and frozen at - 80°C. Samples were thawed on ice and spun for 10 min at 16k × *g* before application of 50 μl to the GE Superdex 200 5/150 column which was equilibrated to SAXS-buffer. SAXS parameters are described in Table S1. Primary data reduction was done in ScatterBrain (2.710), all other data analyses were performed using ATSAS package 2.8.2 or ATSAS online (SASREF and DAMMIN) (Franke et al., 2017). For buffer subtraction, 30 frames before protein elution were selected, averaged and subtracted from the averaged data. Some particle deposition on the capillary was noted in the buffer region impairing the buffer subtraction. Therefore, the first three points were omitted for Guinier analysis. Guinier, distance distribution and Kratky analyses were carried out using Primusqt. The molecular weight of the apoD tetramer was calculated using the Porod volume and *I*(*0*) (Petoukhov et al., 2012), using contrasts and partial specific volumes calculated using MULCh (Whitten et al., 2008).

To create models based on SAXS of the apoD tetramer, 21 conformations from a molecular dynamics simulation of monomeric apoD with modelled glycosylation (Oakley et al., 2012) were used as starting models. Two SASREF runs were performed to model a tetramer against the experimental SAXS data with *D*_*2*_ symmetry (P222 in SASREF). To restrain the model, the four unambiguously inter-subunit BS^3^ crosslinks identified in XL-MS were used with a maximum distance of 32 Å (*B2, B3, B4, B9,* Table 1). The maximum distance was set slightly longer than the theoretical maximum linker distance for BS^3^ of 29 Å to allow for conformational flexibility of sidechains, rearrangement during oligomerisation as well as to prevent the model becoming trapped in a conformation that it consistent with the cross-linking data, but inconsistent with the SAXS data. The 42 produced SASREF models were visually inspected for fulfilling the identified crosslinks, glycosylations interfering with the inter-subunit interface and obvious clashing of glycosylations as SASREF only applies penalties for Cα-Cα clashes (Petoukhov and Svergun, 2005). Two SASREF models were then chosen based on their subunit conformation and according to the evaluation of minimal steric clashing of glycosylations, glycosylation interference with the inter-subunit interface and satisfying the identified crosslinks that were not restrained.

**Table 1.** All crosslinks identified in this study.

| ID | Cross-linker | Residue |  | C-C Distance between residues (Å)<sup>†</sup> |  |  |  |
| --- | --- | --- | --- | --- | --- | --- | --- |
|  |  | 1<sup>st</sup> | 2<sup>nd</sup> | Intra-subunit |  | Inter-subunit |  |
|  |  |  |  | Model 1 | Model 2 | Model 1 | Model 2 |
| *B1* | BS<sup>3</sup> | K21 | K55 | 14.0 | 13.6 | 35.0 | 25.7 |
| *B2* | BS<sup>3</sup> | K55 | K55 | - | - | 27.2 | 25.7 |
| *B3* | BS<sup>3</sup> | K156 | K156 | - | - | 25.5 | 27.0 |
| *B4* | BS<sup>3</sup> | K156 | K155 | - | - | 22.3 | 29.8 |
| *B5* | BS<sup>3</sup> | K167 | K55 | 21.3 | 23.6 | 16.3 | 47.7 |
| *B6* | BS<sup>3</sup> | K167 | K155 | 30.1 | 32.0 | 25.6 | 33.4 |
| *B7* | BS<sup>3</sup> | K155 | K55 | 32.1 | 31.4 | 35.4 | 36.1 |
| *B8* | BS<sup>3</sup> | K144 | K7 | 23.4 | 21.9 | 29.5 | 34.7 |
| *B9* | BS<sup>3</sup> | K144 | K156 | - | - | 14.9 | 16.7 |
| *B10* | BS<sup>3</sup> | K7 | K156 | 20.7 | 20.0 | 34.6 | 23.7 |
| *Z1* | ZLXL | E71 | K55 | 7.4 | 7.5 | 21.9 | 26.8 |
| *Z2\** | ZLXL | K55 | E30 | 22.6 | 22.7 | 32.5 | 38.3 |
| *Z3\** | ZLXL | K155 | E60 | 25.2 | 24.2 | 25.2 | 26.1 |

Note: <sup>†</sup> Distances stated here are as measured in the proposed tetrameric apoD model. \* Not satisfied crosslinks.

In parallel, the experimental SAXS data was used in DAMMIN with P222 symmetry to create 20 bead models in each run. DAMAVER was used for validation and averaging with no model rejected in run 1 and two models rejected in run 2. Using the SUPALM, the filtered bead and SASREF models were aligned and then superimposed in PyMOL. The superimposed SASREF models and the filtered *ab initio* models were visually evaluated for how well they overlay.

### Deglycosylation of apoD

IEX-purified apoD tetramer (0.05 μg) was deglycosylated using PNGase F (glycerol free, P0704L, New England BioLabs) either under denaturing or native conditions according to the manufacturer’s protocol. For denaturing conditions, apoD tetramer was mixed with denaturing buffer, heated to 95°C for 5 min, then glyco buffer, NP-40 and 1 μl PNGase were added and the reaction was incubated for 1 h at 37°C. For native conditions, apoD was mixed with glyco buffer and 1.5 μl PNGase and the reaction was incubated for 15 h, 24 h or 48 h at 37°C. Control samples were treated the same but PNGase was substituted with ultrapure water. The complete sample volumes were analysed on denaturing SDS-PAGE and western blotting as described above, with the difference that protein was transferred to nitrocellulose membranes at 25 V for 90 min and detection was performed on Amersham X-ray films.

## RESULTS

### Characterisation of apoD in three human body fluids

Initial characterisation of apoD in plasma, BCF and CSF was undertaken by SDS-PAGE and western blotting, with sample dilutions optimised for equal signal intensity (Figure 2A). In BCF a single band at approximately 27 kDa was detected, indicative of an apoD monomer. A similar molecular weight was observed for apoD in CSF, while apoD in plasma was slightly larger at approximately 30 kDa. In BCF, a second low abundance band at ∽60 kDa was also immunoreactive to apoD antibody.

**Fig. 2.**
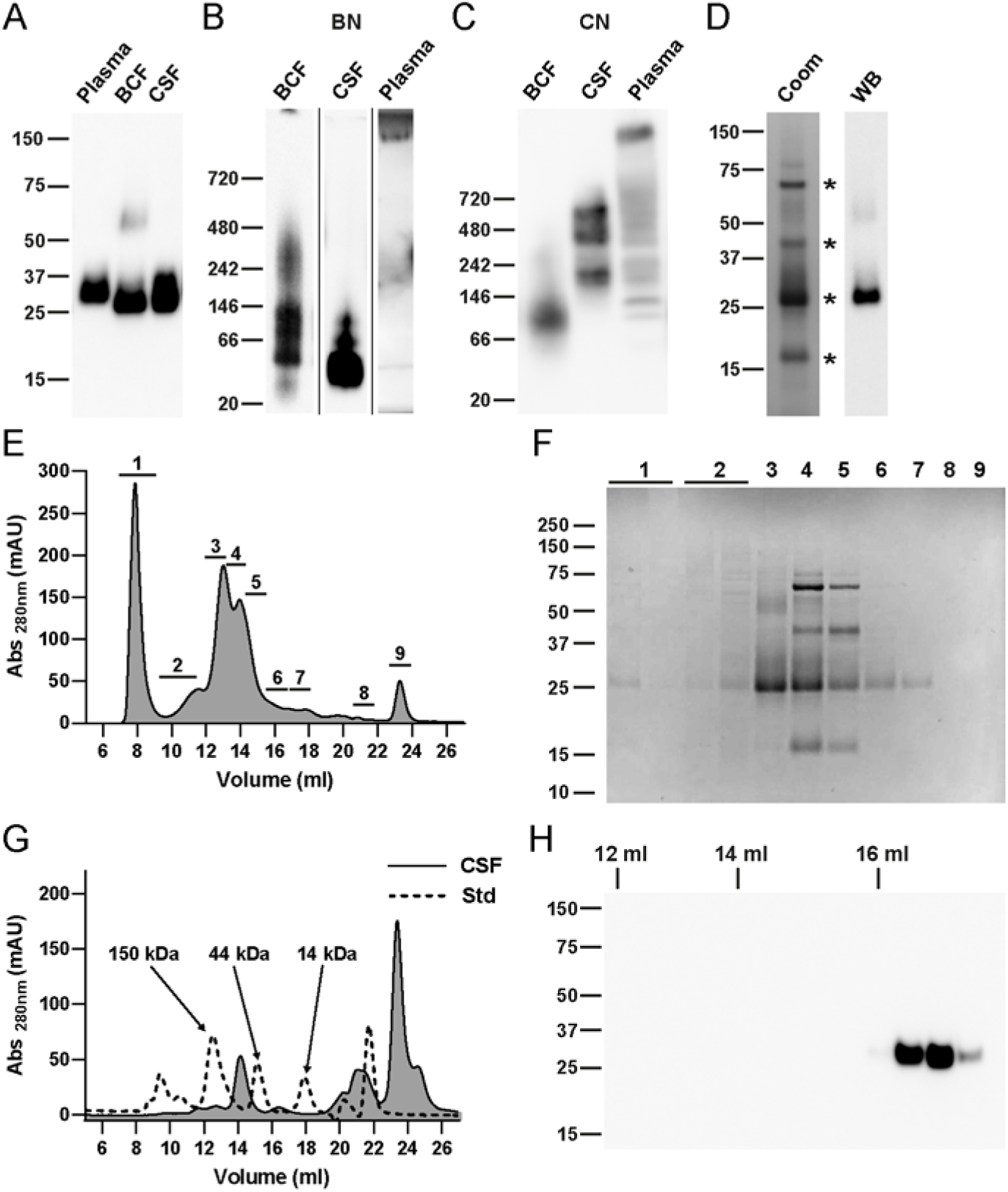
Identification and characterisation of apoD in three human fluids. A) Plasma, BCF and CSF SDS-PAGE western blot probed for apoD showed apoD at ∽27 kDa in BCF and CSF. ApoD in plasma appeared slightly larger. Loading: Plasma: 22 μl of 1:100; BCF: 22 μl of 1:22,000; CSF: 22 μl of 1:15. B) Blue native PAGE (BN) and western blotting of plasma, BCF and CSF revealed higher order apoD species. Loading: Plasma: 10 μl of 1:67; BCF: 10 μl of 1:322.5; CSF: 10 μl of 1:2. C) Clear native PAGE (CN) and western blotting of plasma, BCF and CSF revealed higher order apoD species in BCF. Loading: Plasma: 8 μl of 1:2.29; BCF: 8 μl of 1:533; CSF: 8 μl of 1:4. D) SDS-PAGE Coomassie staining (Coom) of BCF showed the four most abundant proteins (indicated by asterisks) in BCF (albumin, 70 kDa; Zn-α2-glycoprotein, 44 kDa; apoD, 27 kDa; prolactin-inducible protein, 15 kDa). BCF western blot (WB) probed for apoD identified an abundant band at 27 kDa as apoD, as well as a potential dimer species at ∽60 kDa. Loading: Coomassie: 1 μl of 1:10; Western blot: 1 μl of 1:133. E) Size exclusion (Superdex S200 Increase 10/300) UV trace from the direct application of BCF identified nine areas of interest (marked 1-9). F) SDS-PAGE Coomassie of the nine SEC fractions (loading volume 19.5 μl eluate) marked in panel E indicated a high abundance of apoD in peak three. G) Size exclusion (Superdex S200 Increase 10/300) UV trace from the direct application of CSF (unbroken line) and the UV trace of size standards (Std, dashed line). H) SDS-PAGE western blotting (short exposure time) of SEC fractions (loading volume 26 μl eluate) collected in panel G with elution volume indicated. The major apoD portion was detected at 17 ml elution volume.

To assess potential oligomerisation of apoD in BCF, CSF and plasma, samples were analysed using blue native (BN) and clear native (CN) PAGE and western blotting (Figure 2B and C). Using BN PAGE, three major bands were detected in BCF at ∽130 kDa, 70 kDa and 45 kDa. Above and below this molecular weight range, faint smeared bands were also observed. Furthermore, some apoD was present as high molecular weight aggregates in the gel loading well. In contrast, CN PAGE of BCF showed a single band at ∽120 kDa, indicating an oligomeric apoD species. In CSF three bands were identified using BN PAGE, with the main band at ∽30 kDa and additional bands at 60 kDa and 100 kDa. This band pattern was not present in CN PAGE, which showed bands around 600 kDa, 480 kDa and 300 kDa. In plasma, apoD in BN PAGE did not enter the separating gel, indicating a very high molecular mass. Plasma analysed on CN PAGE again showed an apoD band at a similarly high molecular weight, as well as some lower molecular weight apoD bands.

The results of BN PAGE provided a strong indication that apoD is not exclusively present as a monomer in BCF and CSF. Therefore, we analysed apoD oligomerisation in BCF and CSF in greater detail. Four major protein bands were identified in BCF using SDS-PAGE Coomassie staining (Figure 2D, marked with *). According to previous reports, BCF contains albumin (70 kDa), Zn-α2-glycoprotein (44 kDa), apoD (25 kDa) and prolactin-inducible protein (15 kDa) (Balbín et al., 1990; Balbín et al., 1991; Haagensen et al., 1979); it is noteworthy that the major bands observed in the stained gel were consistent with these four proteins. Western blotting again identified apoD as a predominant band at 27 kDa as well as a second low abundance band at 60 kDa (Figure 2D).

To further evaluate apoD oligomerisation in BCF, initial SEC characterisation of BCF was performed directly (<4 hr) after collection through application to a Superdex Increase 200 size exclusion chromatography column (10 mm×300 mm). According to the elution profile depicted in Figure 2E, samples of the main peaks 1-9 were analysed by SDS-PAGE with Coomassie staining (Figure 2F). The predominant protein peak to elute from the column was identified as mainly apoD (peak 3). The retention volume of peak 3 suggests a molecular mass of ∽100 kDa, much larger than the theoretical ∽25 kDa of a monomeric 169 amino acid, fully glycosylated apoD. Notably, bands at the apparent molecular weight of apoB-100 (515 kDa) or apoA-II (11 kDa) were not found at concentrations that are near stoichiometric relevance for apoD. This indicates that the elution volume is not due to an association of apoD with lipoprotein particles. Importantly, identical SEC results were obtained from fresh, non-frozen BCF as from frozen BCF.

Using the same SEC column, SEC analysis was also performed on CSF resulting in the elution profile shown in Figure 2G. No major peak at the exclusion volume (∽8 - 10 ml) was detected, indicating no dominant high-molecular weight complexes, which is in agreement with BN PAGE but not with CN PAGE. SDS-PAGE western blot analysis of fractions within the range of the apoD monomer and oligomers revealed that the major apoD fraction elutes at ∽16.5 ml, corresponding to molecular weight of apoD monomer (Figure 2H). Longer exposure to the CCD camera also showed less abundant apoD at 14.5 ml and 13 ml, which is consistent with the theoretical elution volume of apoD oligomers (Figure S1). The SEC analysis of CSF resembles the BN PAGE analysis but not the CN PAGE, since apoD was not detected in SEC fractions eluting at a corresponding elution volume for proteins larger than 200 kDa (data not shown).

### Size-exclusion characterisation of apoD

To improve chromatographic resolution of the BCF SEC analysis, BCF was next applied to a larger scale 16 mm × 600 mm (16/600) Superdex 200 column, calibrated using size standards (Figure 3A). Fractions over peak 1 and 2 were analysed by SDS-PAGE and probed for total protein by Coomassie staining and for apoD by western blotting (Figure 3B and C). The two main peaks eluted between 60 ml and 80 ml and represented the majority of the protein in BCF. The peak at 66 ml consisted exclusively of apoD at 27 kDa (Figure 3B and C). Importantly, no band stoichiometric to the apoD amount was detected at 11 kDa indicative of apoA-II which apoD would associate with in lipoprotein particles. The second peak at 74 ml was a mixture of all four of the major BCF proteins, including a substantial amount of apoD. Western blot for apoD confirmed the 27 kDa protein as apoD, as well as small amounts of apoD at ∽60 kDa, as observed in western blots of the original BCF prior to SEC analysis (Figure 2). Comparison of the elution profile to the profile of protein standards suggested that the main apoD peak eluted at a volume consistent with an oligomer of ∽120 kDa.

**Fig. 3.**
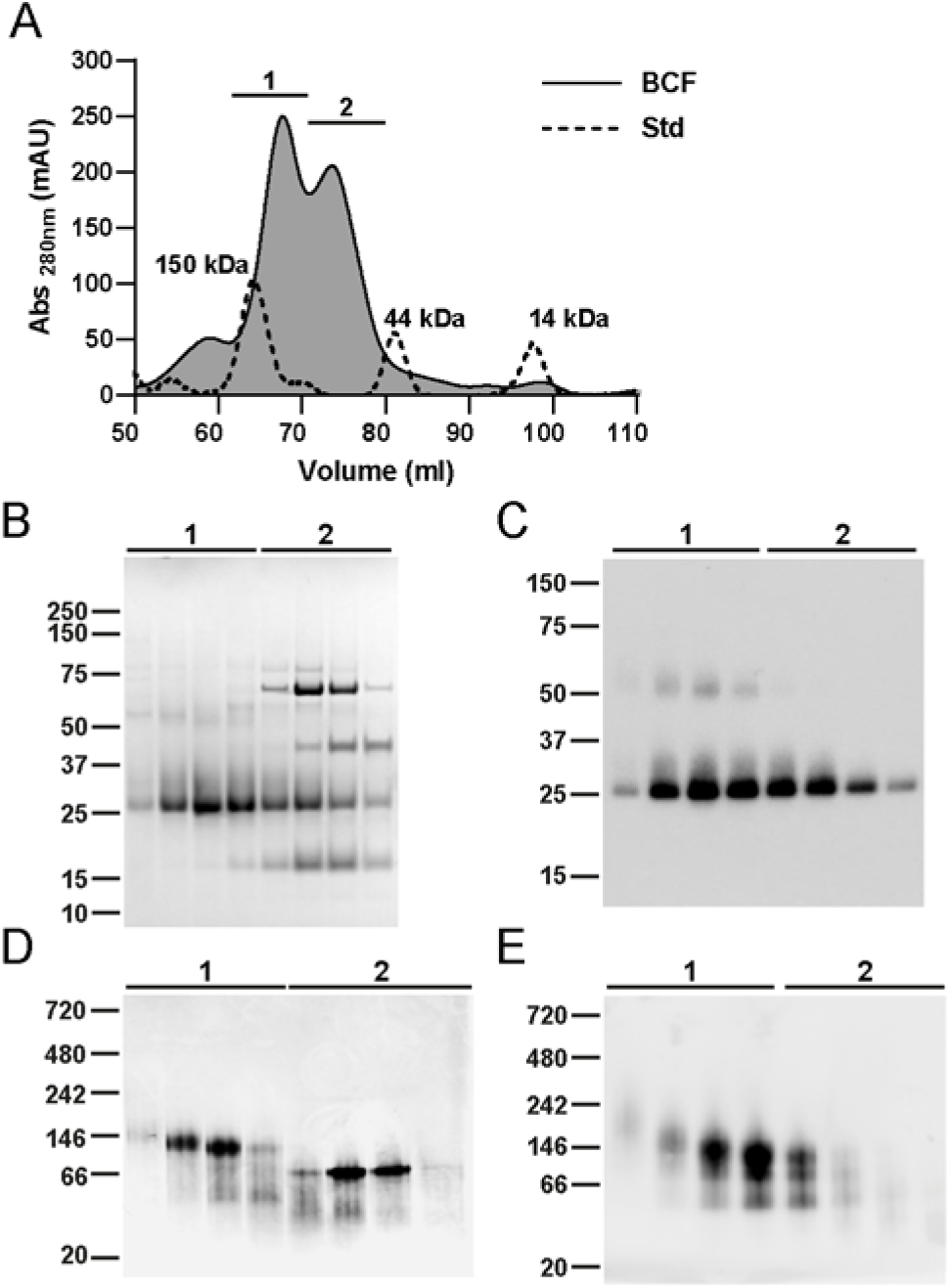
SEC characterisation of apoD in BCF. Neat BCF applied to high-resolution SEC showed that apoD eluted as an oligomer. A) Size exclusion (Superdex S200 16/600) UV trace from the direct application of BCF (unbroken line) and the UV trace of size standards (Std, dashed line). Fractions of 1 ml of eluate over peak 1 from 64.4 ml to 70.4 ml, were pooled and rerun on S200 10/300 (Figure 3). B) SDS-PAGE Coomassie staining (loading volume 19.5 μl eluate) and C) western blotting (loading volume 0.3 μl eluate) of SEC fractions shown in panel A indicated pure apoD in peak 1. D) BN PAGE Coomassie staining (loading volume 30 μl eluate, non-linear adjustment) and E) western blot (loading volume 3.75 μl eluate) of SEC fractions indicated in panel A showed high order apoD species, with the predominant band running at ∽120 kDa.

Results from the large format S200 column confirmed the results obtained from BCF analysis on the small SEC column and from BN PAGE. Specifically, the predominant fraction of apoD eluted from the SEC column at a volume consistent with a hydrodynamic radius at least four-fold larger than expected of monomeric apoD. To validate this result, SEC fractions were then applied to BN PAGE and analysed both by Coomassie staining and western blotting (Figure 2D and E). Coomassie staining (Figure 2D) revealed two major bands: A protein of ∽120 - 140 kDa in peak 1 and ∽70 kDa protein in peak 2. When probed by western blot for apoD (Figure 2E), the 120 kDa band in peak 1 was the predominant band, accompanied by two less abundant bands that migrated further down the gel. The predominant 70 kDa band detected by Coomassie in peak 2 was not immunoreactive with the apoD antibody, which indicates that this band most likely consists of albumin. These results suggest that apoD in BCF predominantly forms a tetramer.

To ascertain if this putative tetrameric apoD species was in dynamic equilibrium with other apoD oligomeric species, S200 fractions containing the ∽120 kDa apoD were pooled, concentrated and reapplied to size exclusion columns in both the 10/300 and 16/600 format. The SEC chromatogram from the 10/300 column is shown in Figure 4A. A single near-symmetric peak was observed with a retention volume consistent with the proposed tetrameric apoD observed in BCF. SEC fractions of the peak were analysed by SDS-PAGE with Coomassie staining. A distinct 27 kDa protein band was detected that was accompanied by a low abundance band at ∽60 kDa (Figure 4B). Both bands were confirmed as apoD by western blotting (Figure 4C). The same fractions were further characterised by BN PAGE, Coomassie staining and western blot (Figures 4D and E). The predominant Coomassie band observed at ∽120 kDa was confirmed as apoD by western blotting. Interestingly, BN PAGE again indicated two smaller protein species below the main 120 kDa band. This may indicate that the apoD tetramer dissociates due to sieving forces applied while migrating through the gel or due to the Coomassie dye acting as a light “detergent” (Wittig and Schagger, 2008).

**Fig. 4.**
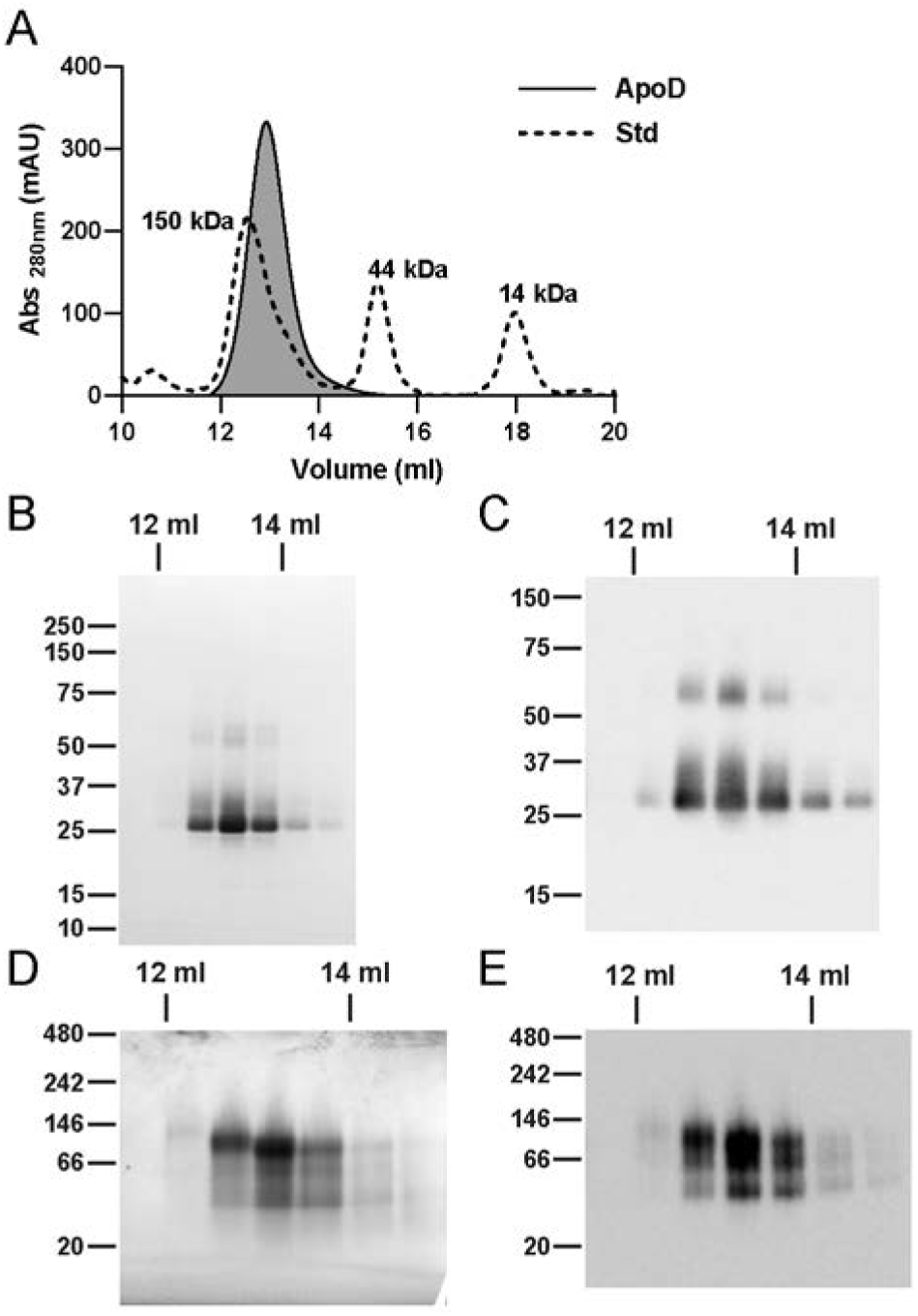
SEC characterisation of apoD purified from BCF. Pooled SEC fractions containing apoD (Figure 2) were rerun on SEC and eluted as a single peak consistent with a hydrodynamic radius of a ∽120 kDa protein. A) Size exclusion (Superdex S200 increase 10/300) UV trace from pooled and rerun apoD fractions (unbroken line) and the UV trace of size standards (Std, dashed line). B) SDS-PAGE Coomassie staining (loading volume 6.7 μl eluate) and C) western blotting (loading volume 0.1 μl eluate) of SEC fractions confirmed that the fractions contained pure apoD. D) Blue Native PAGE Coomassie staining (loading volume 30 μl eluate) and E) western blot (loading volume 1 μl eluate) of SEC fractions showed higher order apoD species, with the predominant band running at ∽120 kDa.

For size determination of the putative apoD tetramer, calibrated SEC columns were used to extrapolate a molecular mass. Retention volumes for both apoD in BCF (Figure 3 peak 3) and purified apoD (Figure 4) from three different SEC column formats were obtained. In addition to the Superdex columns described above, an HPLC-based TOSOH SEC column (∽14 ml CV) was utilised. The HPLC column provided the ability to generate data in triplicate as well as confirm the Superdex SEC data with an alternative chromatographic media. Calibration curves and apoD results are shown in Figure S2. The S200 Increase 10/300 was found to have the best statistical fit within the mass range of the size standards. The calculated apoD size for this column was 124 kDa (BCF apoD) and 131 kDa (purified apoD). Depending on column format and column matrix, the calculated molecular masses of the putative apoD tetramer ranged between 108 and 139 kDa.

Purification of apoD by SEC alone did not yield in sufficient quantity and quality for further structural analyses. Therefore, we used two parallel purification strategies, an established IEX-purification and a novel HIC-purification, each with a subsequent SEC polishing step. To illustrate that these purification strategies did not affect the oligomeric status of apoD, we compare the elution profiles of BCF to SEC-, IEX-and HIC-purified apoD (Figure S3). The SEC elution volume of fresh, not freeze-thawed BCF was consistent with SEC-purified apoD from frozen BCF and HIC-purified apoD (S200 10/300, Figure S3A). Furthermore, the SEC elution volume of BCF was consistent with IEX-purified apoD (S200 16/600, Figure S3B).

### Crosslinking of apoD tetramer

The molecular weight determination of apoD oligomer using BN PAGE and SEC provided useful approximate molecular mass values that were, however, variable to some degree. Therefore, we characterised the apoD tetramer with orthogonal and non-matrix based techniques including protein crosslinking, AUC and MALLS.

Crosslinking of the predominant putative apoD tetramer was performed by crosslinking primary amines using bis(sulfosuccinimidyl)-suberate (BS^3^). The linker arm of BS^3^ is 11.4 Å long when fully extended, allowing a theoretical maximum distance of ∽29 Å between α-carbons of crosslinked residues (Merkley et al., 2014). Crosslinking was performed with both BCF and purified tetrameric apoD for 30 min and 1 h. Crosslinked samples were analysed by SDS-PAGE and western blotting. Figure 5A shows the western blot for crosslinking for 30 min (identical results were obtained after 1 h of crosslinking and are shown in Figure S4). For crosslinking of BCF and purified apoD, an immunoreactive band at 100 - 110 kDa was detected with increasing molar excess of crosslinker. Bands at ∽27 kDa, 50 - 55 kDa and 75 kDa were detected, depending on the BS^3^ concentration. This result indicates that apoD forms a tetramer of ∽100 kDa, which is captured with increasing BS^3^ amounts. Partially crosslinked tetramer dissociates to monomeric, dimeric and trimeric apoD (27, ∽50 and 75 kDa) on SDS-PAGE. Nonspecific crosslinked products (i.e. with other proteins in BCF or very high molecular weight apoD species above the tetramer) were not detected. Quantification of crosslinking western blots showed an increase in apoD tetramer with increasing BS^3^ concentrations for both BCF and purified apoD (Figure 5B and C). This crosslinking experiment confirms the nature of the apoD oligomer as a tetramer.

**Fig. 5.**
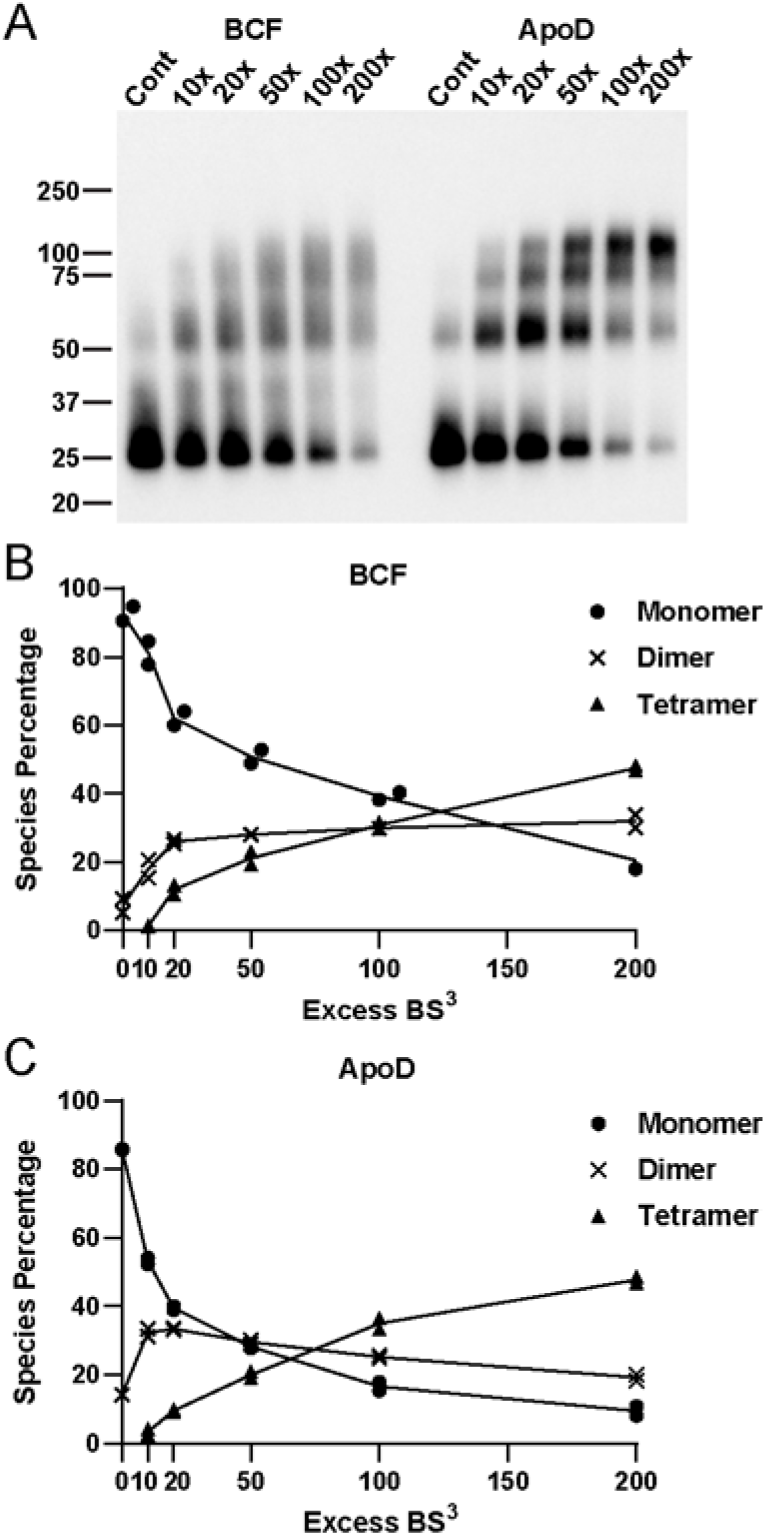
Crosslinking of apoD tetramer. BCF and purified apoD tetramer were crosslinked with BS^3^ using a molar crosslinker excess ranging from 10× to 200×. A) Crosslink reaction mixtures (loading volume 0.13 μl sample of reaction, 0.03 μg apoD) analysed by western blot after 30 min. With increasing crosslinker concentration, fully crosslinked apoD tetramer appeared at ∽100 kDa. No higher size species or aggregates were detected. Cont: Control, no BS_3_. Quantification of crosslinking western blots for B) BCF and C) purified apoD tetramer demonstrated a decrease in monomers and an increase in tetramers corresponding with increasing BS^3^ excess. Percentages of apoD species for 30 min and 1h of crosslinking (blot in Figure S4) are shown at each crosslinker concentration and the black lines connect the means of 30 min and 1 h of crosslinking.

### Multi-angle laser light scattering (MALLS) analysis

For further verification of the apoD tetramer molecular weight, we applied HIC-and IEX-purified apoD tetramer to a Superdex S200 10/300 column combined with MALLS (SEC-MALLS). From SEC-MALLS (Figure 6A) a single, near-symmetrical peak of apoD eluted at a volume corresponding to a tetramer, consistent with the initial SEC characterisation (Figure 3). The observed experimental molecular weight of apoD, weight-averaged from three separate runs, was 93.6 ± 3.7 kDa which is consistent with an apoD tetramer. Interestingly, the MALLS analysis indicated a molecular weight ranging from 110 kDa to 85 kDa over the elution peak and the light scattering signal slightly preceded the UV and differential refractive index measurement (Figure 6A). This shift in the light scattering signal can indicate that the apoD peak may not be entirely homogeneous, e.g. due to varying degrees of apoD glycosylation. This was tested by SDS-PAGE western blot analysis of collected MALLS fractions which demonstrated consistent apparent molecular weight of the monomer units within the apoD tetramer (Figure 6B). Therefore, the apparent heterogeneity of apoD tetramer based on the light scattering profile and the observed range of molecular sizes is not due to differences in gross glycan content.

**Fig. 6.**
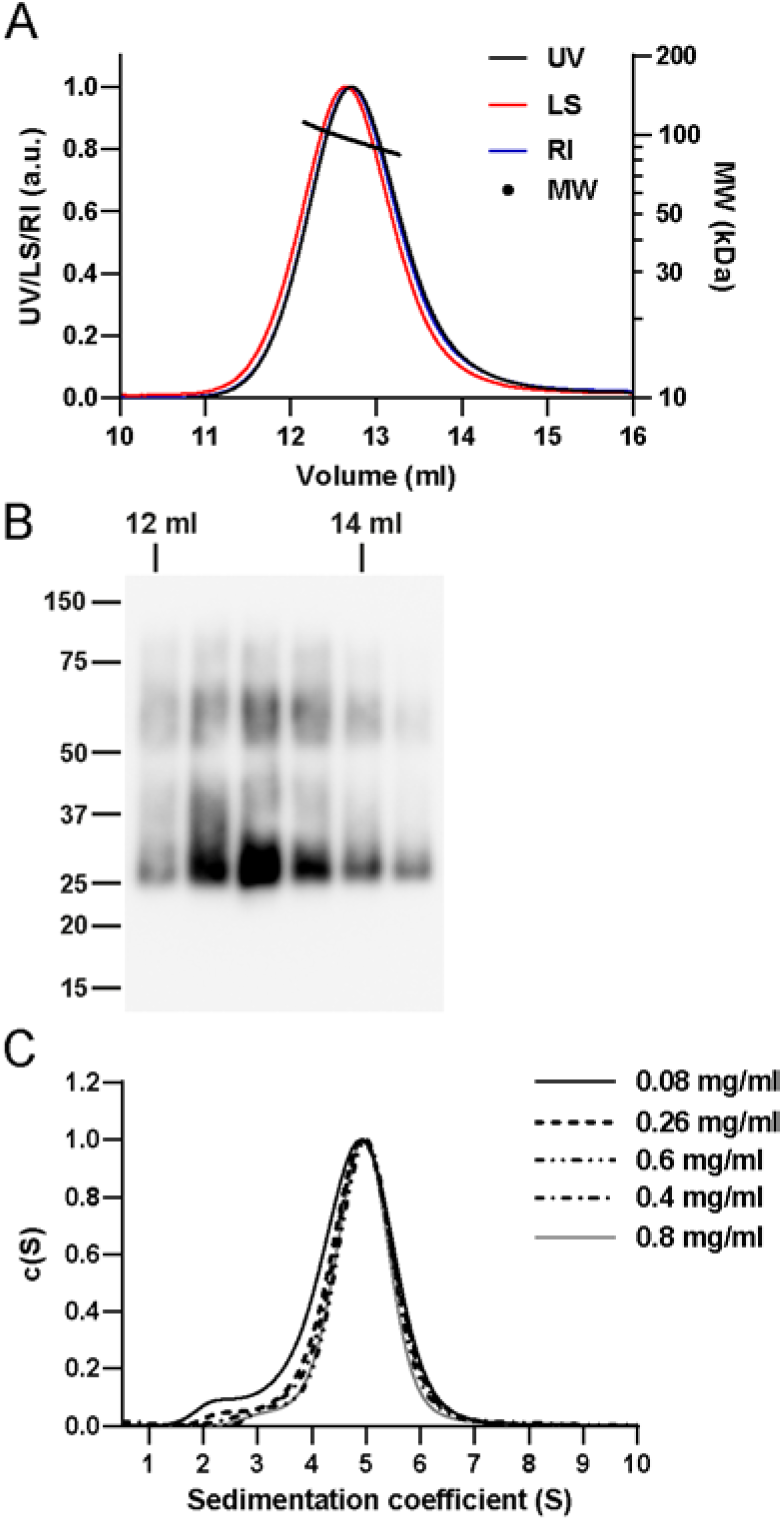
SEC-Multi-angle laser light scattering and analytical ultracentrifugation analysis of apoD tetramer. A) SEC-MALLS was performed on HIC-and IEX-purified apoD tetramer with recordings UV absorbance at 280 nm (UV, black line), light scatter (LS, red line), differential refractive index (RI, blue line) and calculated molecular weight (dots) (representative run). The calculated average molecular weight for the apoD tetramer was 93.6 ± 3.7 kDa (three separate runs) and the molecular weight ranged from 110 kDa to 85 kDa over the peak. The light scattering signal slightly preceded the UV and refractive index signals. B) SDS-PAGE Coomassie staining of eluted fractions (loading volume 26 μl eluate) revealed symmetrical apoD bands consistently at 27 kDa. C) Sedimentation velocity AUC was carried out with purified apoD tetramer at indicated concentrations and the distribution of sedimentation coefficients was determined. ApoD tetramer had a sedimentation coefficient of ∽4.9 S and was relatively stable upon dilution. At a concentration of 0.08 mg/ml, small amounts of apoD monomer with a sedimentation coefficient of ∽2.4 S could be detected.

### Analytical ultracentrifugation analysis of apoD tetramer

Sedimentation velocity by analytical ultracentrifugation (AUC) was utilised to further confirm the oligomeric state of apoD and identify a potential dynamic equilibrium between different oligomeric species upon dilution. The extensive glycosylation of apoD made precise determination of molecular weight by sedimentation equilibrium unfeasible as detailed below.

Sedimentation velocity of tetrameric apoD was measured at a range of apoD concentrations from 0.08 mg/ml to 0.8 mg/ml and the normalised distribution of sedimentation coefficients is shown in Figure 6C. ApoD tetramer showed a predominant peak at a weight-average sedimentation coefficient of 4.9 ± 0.630S. This corresponds to a molecular weight of roughly 85 kDa assuming a globular, non-glycosylated protein. The broadness of the apoD peak and the deviation in molecular mass (as compared to the SEC and PAGE techniques described above) may be due to the extensive glycosylation of apoD that makes the protein more buoyant (Lebowitz et al., 2002). Upon dilution, apoD tetramer remained stable, however, when apoD concentration was decreased to 0.08 mg/ml, a small fraction (∽6%) of apoD was detected at a sedimentation coefficient of 2.4 S. This agreed with the sedimentation coefficient of HIC-purified apoD monomer (data not shown), corresponding to a molecular weight of around 26.5 kDa, matching an apoD monomer.

### SEC-Small-angle X-ray scattering (SEC-SAXS)

Considering our results indicating a novel native tetrameric form of apoD, size-exclusion coupled with small angle X-ray scattering (SEC-SAXS) was applied to purified tetrameric apoD. SEC was utilised to remove traces of aggregates in-line prior to SAXS analysis. Freeze-thawing and glycerol in the SEC buffer to avoid radiation damage were shown by SEC to not affect the oligomeric status of apoD (Figure S3). All SAXS experimental and calculated parameters are presented in Table S1. Post-collection analysis of the radius of gyration (*R*_*g*_) and scattering intensity *I*(*0*) over the SEC peak determined the region of constant *R*_*g*_ for averaging and further analysis (marked in red, Figure 7A). After averaging and buffer subtraction, the data was evaluated in the Guinier region (Figure 7B). The data at small *q* values did not show any up-or down-turn. Together with the *R*_*g*_ stability over time in the selected, averaged region, this indicates no aggregation or interparticle repulsion. The SAXS intensity profiles *I(q) versus q* for experimental values (yellow) and two rigid body models (SASREF model 1, orange and 2, cyan) are shown in Figure 7C. All experimental and calculated parameters are listed in Table S1. For the experimental values, the *I(q) versus q* profile was transformed to a *P*(*r*) function shown in Figure 7D. *P*(*r*) corresponds to the probable distribution of vector lengths (*r*) between scattering centres within the scattering molecule and can be used to determine approximate shape and size of scattering molecules. The *P*(*r*) profile of tetrameric apoD corresponded to a globular shaped molecule, with a Porod volume of 169000 Å^3^. Based on the Porod volume, apoD molecular weight was estimated to be 99 kDa. This is in good agreement with molecular weight determination based on *I*(*0*) which is 97.1 kDa. Therefore, SEC-SAXS of oligomeric apoD further supports the tetrameric state of apoD. Additionally, the Kratky plot in Figure S5A showed a bell-shaped curve indicating that the apoD tetramer has a globular shape.

**Fig. 7.**
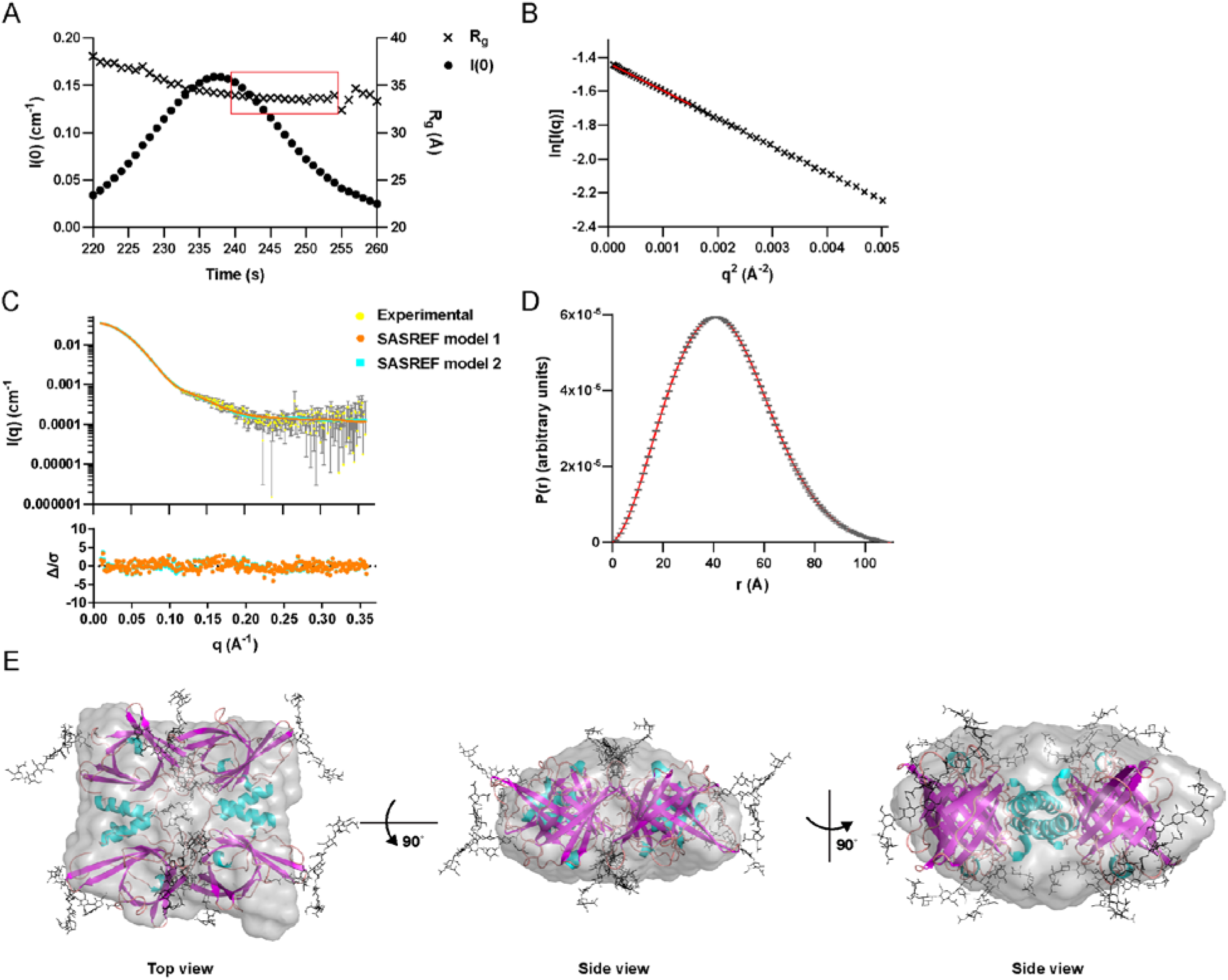
SEC-SAXS analysis of apoD tetramer. A) Scattering intensity I(0) (dots) and radius of gyration R_g_ (crosses) over elution of apoD on the S200 5/150 SEC column identified the region for averaging (marked in red). B) Guinier plot of apoD tetramer and linear regression, indicating no up-or down-turn of the curve. C) Experimental scattering profile of apoD tetramer (yellow dots with grey error bars). Continuous lines represent the calculated scattering profiles of the two SASREF rigid-body models shown in panel D (model 1, orange rhomb, χ^2^=1.25) and Figure S5 C (model 2, cyan square, χ^2^=1.44). The lower inset plot shows the error-weighted residual Δ/σ difference plot. D) P(r) function of apoD tetramer. The symmetric profile was indicative of a mainly globular scattering molecule. In panels C and D, errors are based on counting statistics and error bars are not shown if they are smaller than symbol sizes. E) Representative model of apoD tetramer generated by the combination of ab initio model 2 (grey) and rigid-body model 1 (cartoon showing β-sheets in pink, α-helices in cyan and loops in tan, with glycosylations in black) is depicted in top and side views. The entry to the ligand binding site was facing outwards and seems accessible. The inter-subunit interface was comprised of stacked adjacent α-helices and some parts of the β-barrel and their loops. The two models overlaid reasonably well, with some glycosylations, the end of two β-sheets and their loops protruding.

### Chemical crosslinking and mass spectrometry (XL-MS)

To gain more insight into the tetrameric structure of apoD, we utilised XL-MS to obtain distance information. Purified apoD was crosslinked with either BS^3^ (crosslinks primary amines), ADH (crosslinks carboxylic acids) or DMTMM, a side-reaction in the ADH crosslinking experiment, which crosslinks primary amines to carboxylic acids. DMTMM catalysed reactions will be referred to as ‘ZLXL’ (zero-length crosslinks) from here on. The use of crosslinkers with varying reactivities serves to enhance the dataset by providing complementary orthogonal data. Briefly, in the XL-MS workflow, the protein samples were crosslinked, digested to peptides, then crosslinked peptides were enriched via SEC, and finally analysed by high-resolution LC-MS/MS. Crosslinked peptides were identified from the MS/MS data by pLINK software (Yang et al., 2012). Only crosslinks that were unambiguously identified with high confidence were used in subsequent analyses (see Materials and Methods for conditions). From two purified samples (6 crosslinking reactions), a total of 13 unique, unambiguous and high-confidence crosslinks were obtained. We observed 10 crosslinks from the BS^3^ experiments and 3 crosslinks from the ZLXL experiments. No crosslinks were identified using the ADH crosslinker. The identified crosslinks are listed in Table 1 and their respective MS spectra are shown in Figure S6.

Out of these 13 crosslinks, four crosslinks (Table 1, crosslinks *B2, B3, B4* and *B9*) were found to be unambiguously inter-subunit. This was based on the observation that the crosslink was either identified between identical residues (*B2*: K55-K55 and *B3*: K156-K156) or between residues derived from peptides with overlapping sequences (*B4*: peptide 1 (residues 156-169) crosslinked to peptide 2 (residues 145-156), and *B9*: peptide 1 (residues 132-156) crosslinked to peptide 2 (residues 156-167)). For these crosslinks to occur, the peptides must have originated from two apoD monomers. In addition, three crosslinks (Table 1, crosslinks *B7, Z2* and *Z3*) were also noted to be possibly inter-subunit crosslinks rather than intra-subunit; the measured intra-subunit crosslinker lengths for these three crosslinks exceed the theoretical length of the crosslinker.

### SEC-SAXS modelling

To further our structural understanding of tetrameric apoD, we applied an *ab initio* modelling approach using DAMMIN to produce low-resolution shapes with P222 symmetry for the tetrameric structure based on the SAXS data. The two filtered models are shown in Figure S5 B and exhibit no major differences in shape (model 1, orange: Normalised spatial discrepancy (NSD)=1.315 ± 0.077; model 2, cyan: NSD=1.203 ± 0.105).

In parallel, monomeric models of apoD high-resolution X-ray structure (Eichinger et al., 2007) with different modelled glycosylation conformations (Oakley et al., 2012) were used as rigid bodies for SASREF modelling. We assumed a D_2_ symmetry since D_2_ is a very common symmetry for a tetramer (Goodsell and Olson, 2000). A C_4_ symmetry is another possible symmetry for a tetramer, however, models with C_4_ symmetry had a worse fit with the experimental scattering data (χ^2^ > 2.1, data not shown). We chose the unambiguously inter-subunit BS^3^ crosslinks *B2, B3, B4* and *B9* as restraints in the modelling of the apoD tetramer, as BS^3^ crosslinking was also experimentally tested to not introduce aggregation of apoD (Figure 5). Two rounds of SASREF modelling using all 21 glycosylation conformations (Oakley et al., 2012) were performed and the resulting models were visually evaluated and selected based on minimal steric clashing of glycosylations, glycosylation interference with the inter-subunit interface and satisfying the identified crosslinks that were not restrained.

Two global subunit conformations resulted from the modelling process. One representative model for each conformation was selected (SASBDB accession number: SASDD83) and the theoretical scattering is shown in Figure 7C (model 1: orange, model 2: cyan). The theoretical scattering of both models corresponds well with the experimentally determined scattering (model 1: χ^2^=1.28, model 2: χ^2^=1.26). This agreement is also highlighted by the error-weighted residual difference plots (I_exp_(q) - I_*calc*_(q))/σ_*exp*_(q) vs q) in the lower panel in Figure 7C: The residual plot shows a flat curve without pronounced deviation, including in the high q area.

Finally, the rigid-body tetramer models were structurally aligned with *ab initio* model 2 which produced a better overlay than *ab initio* model 1. The resulting overlay models for the apoD tetramer are shown in Figures 7E and S5C in top and side views. SASREF models are depicted as a cartoon showing β-sheets in pink, α-helices in cyan and loops in tan, with glycosylations in black, the *ab initio* model is shown in a grey surface representation. The distances measured in the models for all XL-MS identified crosslinks can be found in Table 1. Both models fulfilled all BS^3^ crosslinks and the inter-subunit BS^3^ crosslinks were identified between symmetric and asymmetric subunits. This fits the finding of a partially crosslinked apoD tetramer which dissociated to a trimer on SDS-PAGE (Figure 5).

In SASREF model 1, the ligand binding pocket of apoD was facing outwards and accessible to ligands (Figure 7E). This conformation was obtained in approximately 75% of all models. The tetramer was built of antiparallel stacked β-barrels and stacked adjacent α-helices. The inter-subunit interface of model 1 consisted of stacked adjacent α-helices and some parts of the β-barrel and their loops. The *ab initio* and rigid-body model overlaid well (NSD=1.6813). Minor deviations were found in flexible regions such as glycosylations, loops and parts of β-sheets. In comparison, in SASREF model 2, the orientation of the ligand binding pocket was substantially different (Figure S5C). The β-barrels of the subunits were facing towards each other and appear inaccessible to ligands. The subunit-interface consisted of loops between β-sheets. Again, the *ab initio* and rigid-body model overlaid well (NSD=1.7155). Minor deviations were found in glycosylations, loops and parts of β-sheets. Interestingly, in both models, the four redox-active M93 side chains were fairly exposed and accessible. This is consistent with their previously described antioxidant activity that is due to interaction with L-OOHs (Bhatia et al., 2012b).

## DISCUSSION

Oligomerisation is a common feature in the lipocalin family that can influence ligand binding behaviour in some lipocalins (Gamiz-Hernandez et al., 2015; Gutierrez-Magdaleno et al., 2013). ApoD, however, has been generally considered as a monomer (Akerstrom et al., 2000) but is known to dimerise upon reducing peroxidised lipids (Bhatia et al., 2012a). Here, we show that apoD is mainly present as a tetramer in BCF and that apoD oligomerisation may also take place in CSF.

ApoD was detected at slightly different apparent molecular weights in plasma, BCF and CSF on SDS-PAGE western blots (Figure 2A). This is most likely the result of differential apoD glycosylation, which is known to depend on the tissue location of apoD expression (Li et al., 2016; Schindler et al., 1995; Zeng et al., 1996). BN PAGE revealed oligomeric apoD species in BCF and CSF, whereas in plasma, apoD was detected at a very high molecular weight, as confirmed by CN PAGE (Figure 2). This is consistent with the previously described association of apoD with HDL particles in plasma (Blanco-Vaca et al., 1992). In CSF, BN PAGE and SEC indicated the presence of oligomeric apoD, whereas with CN PAGE, apoD was found in higher molecular weight apoD bands (Figure 2). This suggests that apoD monomer and oligomer may be associated with small lipoprotein particles in CSF and that this association dissipates upon BN PAGE and with SEC. ApoD has been previously shown to associate with lipoproteins in CSF, more specifically with particles containing apoE, apoA-I and-IV, apoJ and apoH, and with larger particles containing apoE, apo-IV and apoJ (Koch et al., 2001). In this previous publication (Koch et al., 2001), SEC on CSF was performed and apoD was shown to co-elute with apoE in the major lipoprotein peak, as well as to elute later, together with apoA-IV. Unfortunately, elution volumes and calibrations were not reported. In the present study we used non-concentrated CSF, whereas Koch and co-workers used 100-fold concentrated CSF (Koch et al., 2001). These differences in purification and concentration techniques applied could account for differences in apoD oligomeric states detected.

In BCF, the main apoD species detected by native PAGE and SEC was an apoD oligomer of around 120 kDa (Figures 2-4). Whereas HDL, LDL and VLDL have been shown to be present in BCF (Mannello et al., 1996; Martinez et al., 1994), we did not find stoichiometric relevant amounts of apoA-II or apoB-100 in the relevant SEC fractions (Figure 2-4). Furthermore, neither apoB-100 nor apoA-II were identified using shotgun LC-MS/MS which was performed in conjunction with XL-MS. This indicates that the elution volume of 120 kDa is not due to association of apoD with lipoprotein particles but a result of oligomerisation. BCF is a fluid produced by benign cysts of the breast which are usually not cancerous (Mannello et al., 2006). Therefore, apoD oligomerisation most likely is not a result of tumourigenesis in breast cysts and may also occur under normal healthy conditions in CSF.

Using SEC and native PAGE to determine the molecular weight of oligomeric apoD, we noticed that these methods created some degree of ambiguity. Size determination on SEC showed variability depending on the column format and column media (Figure S2). This made identification of the exact composition of the apoD oligomer difficult. The observed variation may be due to the extensive glycosylation of apoD, which leads to a different retention of apoD on matrix-based SEC and PAGE compared with non-glycosylated size standards (Andrews, 1965). Indeed, sialylated proteins are known to elute earlier from SEC columns than expected for their actual molecular weight (Alhadeff, 1978). Native PAGE proved to be a valuable tool to assess oligomerisation in fluids with low apoD concentrations, such as plasma and CSF but was not precise enough to detect the exact oligomeric status of apoD. This can be due to the fact that in clear native PAGE retention is influenced by charge in addition to molecular weight. Interestingly, purified apoD tetramer on BN PAGE resulted in three bands (Figures 3 and 4, D and E), indicating that the tetramer dissociated into dimers and monomers. This behaviour could be explained by Coomassie dye acting as a mild “detergent”, which can cause complexes to dissociate. Coomassie especially covers hydrophobic areas (Wittig and Schagger, 2008), of which apoD contains several in extended loops (Eichinger et al., 2007). Our experiments investigating the oligomerisation of a native, glycosylated protein underline the importance of using several complementary methods for studying the molecular weight of protein complexes.

To extend the data derived from SEC and native PAGE, we used protein crosslinking, MALLS and AUC as non-matrix based techniques for molecular weight determination of the apoD oligomer (Figures 5 and 6). All three techniques showed that the apoD oligomer is in fact a tetramer of ∽95 - 100 kDa. This is remarkable since apoD has so far been considered monomeric and to dimerise upon oxidation of M93. Our data show for the first time that apoD displays oligomerisation behaviour similar to other lipocalins. Three proteins that are apoD homologues, or share sequence homology with apoD, form oligomers. Lazarillo, the apoD homologue in the grasshopper *Schistocerca Americana* forms oligomers (30% sequence identity), as shown by SEC (Sanchez et al., 2008). Bilin-binding protein from the butterfly *Pieris brassicae* forms a tetramer, as shown by X-ray crystallography (Huber et al., 1987b). Sandercyanin, an apoD homologue in the North American fish *Sander vitreus* (42% sequence identity), was shown by X-ray crystallography to form a tetramer upon ligand binding (Ghosh et al., 2016). In the light of the prevalence of oligomerisation in the lipocalin family and the uncertainty of the oligomeric status of some lipocalins (Akerstrom et al., 2000), the case of apoD emphasises the usefulness of reviewing the potential oligomerisation of other lipocalins.

Furthermore, controversies over the oligomeric status of certain lipocalins have been attributed to differences in methodological approaches and recombinant cloning strategies (Gasymov et al., 2007; Schiefner et al., 2010). It is noteworthy that the monomeric recombinant apoD used for crystallisation was not glycosylated and contained seven point mutations to allow purification and crystallisation (Nasreen et al., 2006). This discrepancy between the oligomeric state of recombinant and native protein underscores the value of using native protein and employing multipronged approaches when studying protein structure and oligomerisation. Our experiments also highlight the importance of the apoD glycosylation which, intriguingly, depends on the tissue location of expression (Li et al., 2016; Schindler et al., 1995; Zeng et al., 1996). We were unable to fully deglycosylate the apoD tetramer under native conditions even after 48 h (Figure S7). Additionally, the mutated side chains were not located in the ligand binding pocket but on exposed hydrophobic patches (I118S, L120S, C116S) or on the exposed bottom of pocket (L23P) (Eichinger et al., 2007). These changes on the surface may therefore influence self-association of apoD. Whereas BN PAGE and SEC-MALLS pointed to a potential dissociation of the apoD tetramer, apoD seemed fairly stable upon ten-fold dilution in AUC, with only slight dissociation detected. This potentially indicates a low *K*_*D*_ value for the apoD tetramer in a concentration-dependent, dynamic equilibrium. Assessing this question further proved problematic due to detection limits of AUC, MALLS and SEC (absorbance measurements at 214 or 280 nm).

In other lipocalins, ligand binding (Gutierrez-Magdaleno et al., 2013) or pH (Mans and Neitz, 2004; Qin et al., 1998) have been shown to influence oligomerisation. Since ligand binding of apoD may play a role in inflammation and oxidative stress and, therefore, in AD due to binding to AA and other lipids (Bhatia et al., 2013; Phillis et al., 2006), apoD oligomerisation could potentially influence these functions and lead to differentiated behaviour of apoD. Recently, apoD was shown to protect vulnerable lysosomes from oxidative stress and was found to be located on the inside of lysosomes, where pH is low, and to maintain ligand binding (Pascua-Maestro et al., 2017). Therefore, it could be interesting in future to assess the oligomeric structure of intralysosomal apoD.

Our SAXS experiments further confirmed the presence of an apoD tetramer, with a calculated molecular weight of 99 kDa and proposed a globular structure (Figure 7 and S5). Additionally, we provide structural insight into the apoD tetramer. ApoD model 1 showed stacked antiparallel β-barrels and stacked adjacent α-helices and an accessible ligand pocket. In contrast, model 2 showed a different conformation, present in 25% of all models, with ligand pockets facing either each other (Figure S5). These orientations would influence the ligand binding function of the apoD tetramer substantially. Past experiments that used similar purification techniques for apoD demonstrated ligand binding behaviour of apoD from BCF (Ruiz et al., 2014). SAXS however, as a low-resolution technique, cannot definitely determine which subunit orientation is predominantly or exclusively present in the tetramer. Interestingly, in both models, the redox-active residue M93 and glycosylations are accessible for redox activity and recognition, respectively. Furthermore, the so-called ‘spike’, located at the bottom of the apoD ligand binding pocket, is exposed in both models. This area has been implicated in binding to basiginin, a potential receptor for apoD internalisation (Najyb et al., 2015).

In summary, our data revealed that apoD predominantly forms tetramers in BCF and that apoD oligomerisation also takes place in CSF. Our results may have implications for apoD ligand binding and antioxidant function in different tissues and further highlight the oligomerisation propensity of the lipocalin family.

## ACKNOWLEDGEMENTS

Thanks to Kerry-Ann Rye for advice with protein crosslinking, to Timothy Ryan from the SAXS/WAXS beamline at the Australian Synchrotron, part of ANSTO, for help and advice with the SAXS experiments and analysis and to Ann Turnley for proof-reading. BG holds a fellowship from the NHMRC (FT110100249). Travel support for the SAXS experiments was granted by the Australian Synchrotron.

## Abbreviations

AA: Arachidonic acid
AD: Alzheimer’s disease
ADH: Adipic acid dihydrazide
ApoD: Apolipoprotein-D
AUC: Analytical ultracentrifugation
BCF: Breast cyst fluid
BSA: Bovine serum albumin
CSF: Cerebrospinal fluid
CV: Column volume
DMTMM: 4-(4,6-dimethoxy-1,3,5-triazin-2-yl)-4-methylmorpholinium chloride
FDR: False discovery rates
HIC: Hydrophobic interaction chromatography
IEX: Ion exchange chromatography
LC-MS/MS: Liquid chromatography-Tandem mass spectrometry
MALLS: Multi-angle laser light scattering
NSD: Normalised spatial discrepancy
SAXS: Small-angle X-ray scattering
SEC: Size exclusion chromatography
XL-MS: Crosslinking mass spectrometry
ZLXL: Zero-length crosslinks

